# Interactions between terminal ribosomal RNA helices stabilize the *Escherichia coli* large ribosomal subunit

**DOI:** 10.1101/2023.01.19.524821

**Authors:** Amos J. Nissley, Tammam S. Kamal, Jamie H. D. Cate

## Abstract

The ribosome is a large ribonucleoprotein assembly that uses diverse and complex molecular interactions to maintain proper folding. *In vivo* assembled ribosomes have been isolated using MS2-tags installed in either the 16S or 23S ribosomal RNAs (rRNAs), to enable studies of ribosome structure and function *in vitro*. RNA tags in the *Escherichia coli* 50S subunit have commonly been inserted into an extended helix H98 in 23S rRNA, as this addition does not affect cellular growth or *in vitro* ribosome activity. Here, we find that *E. coli* 50S subunits with MS2 tags inserted in H98 are destabilized compared to wild type (WT) 50S subunits. We identify the loss of RNA-RNA tertiary contacts that bridge helices H1, H94, and H98 as the cause of destabilization. Using cryogenic electron microscopy (cryo-EM), we show that this interaction is disrupted by the addition of the MS2 tag and can be restored through the insertion of a single adenosine in the extended H98 helix. This work establishes ways to improve MS2 tags in the 50S subunit that maintain ribosome stability and investigates a complex RNA tertiary structure that may be important for stability in various bacterial ribosomes.

## INTRODUCTION

The ribosome is a large ribonucleoprotein assembly responsible for protein synthesis in the cell. Due to its complexity, the ribosome is composed of intricate networks of RNA-protein and RNA-RNA interactions (Noller 2005). These complex interactions are formed during ribosome assembly, which is a process requiring ribosomal RNA (rRNA), ribosomal proteins (rProteins), and many auxiliary assembly factors (Shajani et al. 2011). In *Escherichia coli* (*E. coli*), the three rRNAs–5S, 16S, and 23S–are all transcribed as a single transcript. During the early stages of ribosomal large subunit (LSU) assembly, sequences flanking the 5′ and 3′ ends of the pre-23S rRNA transcript base-pair to form the leader trailer (LT) helix (Liiv and Remme 1998). The LT helix is then subsequently cleaved by several RNases, and the remaining RNA helix comprises helix 1 (H1) of the pre-23S rRNA (Ginsburg and Steitz 1975; Nikolaev et al. 1973; Li et al. 1999). Additionally, it has been shown that some bacteria remove H1 post-assembly and this removal is correlated with the evolutionary loss of H98 (Shatoff et al. 2021), which suggests a synergistic role between the two helices.

The use of affinity tags in the 23S rRNA has enabled the *in vitro* study of *in vivo*-assembled LSUs with mutations in 23S rRNA. This has commonly been achieved through the insertion of an RNA helix from the genome of the MS2 bacteriophage into helix H98 of the 23S rRNA on the surface of the ribosome (Youngman and Green 2005). Tagged mutant ribosomal subunits can then be purified using the MS2 coat protein dimer that selectively binds to the MS2 RNA helix (Peabody 1993). MS2 tags inserted in H98 of 23S rRNA have been used extensively to enable the investigation of rRNA function (Youngman et al. 2004; Lancaster et al. 2008), in engineered ribosomes (Ward et al. 2019), and for *in vivo* ribosome tracking (Metelev et al. 2022). Although *E. coli* 50S ribosomal subunits with MS2 tags inserted into H98 retain wild-type (WT) *in vivo* and *in vitro* function, we previously showed that these 50S subunits have lower thermostability compared to WT 50S subunits (Nissley et al. 2023).

Here we investigate the loss of thermostability in the *E. coli* LSU upon addition of an MS2 tag into H98. We find that RNA contacts between nucleotides at the interface of rRNA helices H1, H94, and H98 are important for the stability of the *E. coli* 50S ribosomal subunit. Using cryo-EM, we show that this interaction is disrupted by the addition of previous MS2 tag designs in H98 and that the addition of a single adenosine residue in the tag can restore the tertiary interaction and WT levels of stability. The MS2-tags described herein offer an improved method for studying mutant and engineered ribosomes without the introduction of additional instability. Moreover, tertiary interactions between helices at the terminal ends of 23S rRNA corresponding to H1, H94, and H98 are found in many bacteria and these likely play stabilizing roles in a variety of bacterial ribosomes.

## RESULTS

### H1, H94, and H98 interact in the *E. coli* LSU

To better understand the architecture of the region surrounding H98, we examined the structure of the *E. coli* ribosome (Watson et al. 2020). Helices H1, H94, and H98 reside on the surface of the 50S ribosomal subunit, and the apical regions of H1 and H98 extend away from the ribosome (**Figure 1A**). H1 is formed through base pairing between the 5′ and 3′ ends of the 23S rRNA (nucleotides 1-8 and 2895-2904 respectively). The 3′ end of 23S rRNA comprises Domain VI of the LSU which contains RNA helices H94-H101. In the *E. coli* ribosome, H1, H94, and H98 are in close proximity (**Figure 1B**), and nucleotides at the base of H1 (G9, A10), the base of H94 (U2629), and in the apical loop of H98 (A2800) form tertiary interactions.

**Figure 1.**
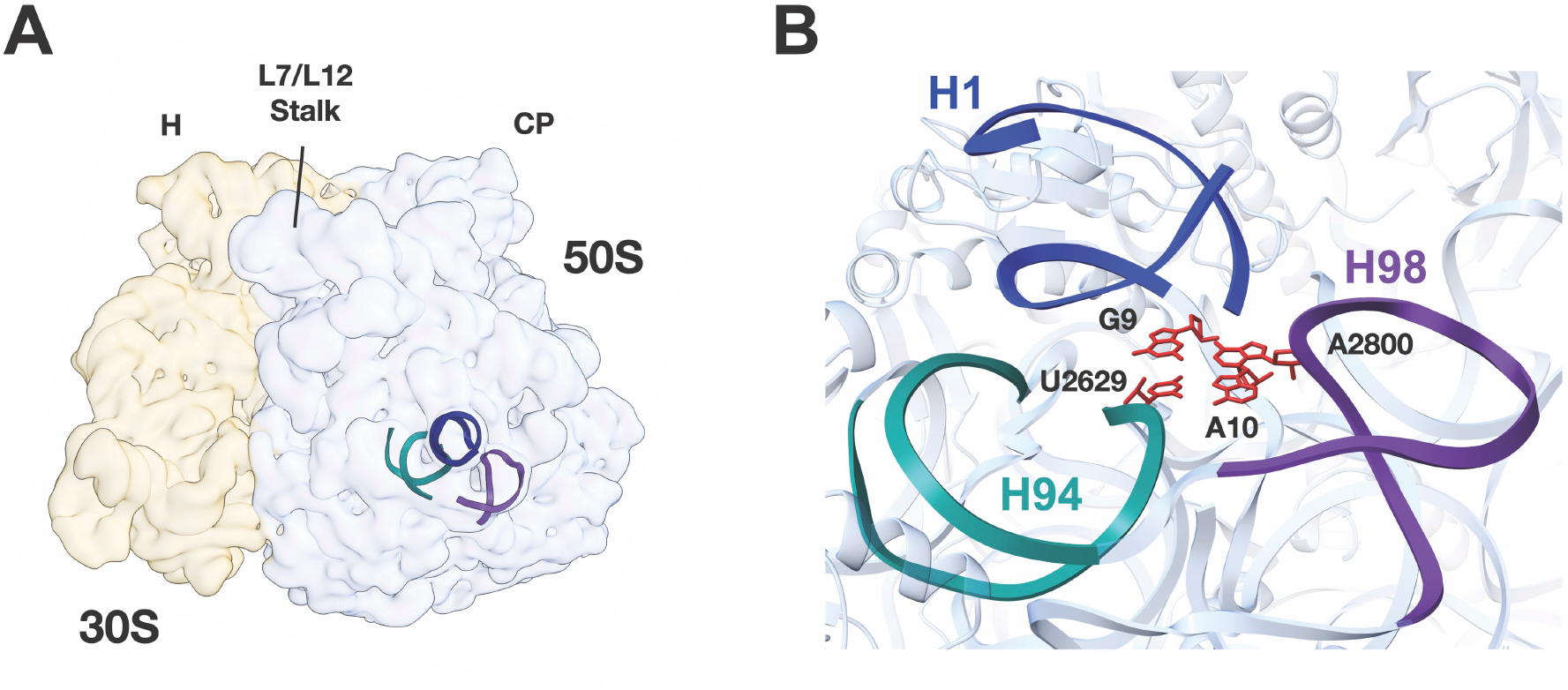
Location and overview of the region surrounding 23S rRNA helix H1. A) Location of helices H1 (dark blue), H94 (teal), and H98 (purple) in the *E. coli* LSU. The locations of the Central Protuberance of the 50S subunit (CP) and Head of the 30S subunit (H) are marked. B) Structural model of the *E. coli* 70S ribosome (PDB:7K00) highlighting interactions between H1, H94, and H98. Nucleotides that form the interaction between these helices are colored red.

The purification of *E. coli* LSUs with mutations in 23S rRNA has commonly been achieved through the insertion of MS2 tags into H98. The original sequence of the MS2 tag from (Youngman and Green 2005) is shown in **Figure 2A** (referred to here as MS2-V1). The tag is inserted into H98 by extension of the helix, which culminates with a poly-uridine stretch (RNA linker). The MS2 helix is then inserted at the end of the RNA linker to ensure that it is accessible to the MS2 coat protein. A second tag design, which includes a different linker sequence and mutations in the MS2 stem loop that increase the MS2 coat protein’s affinity for the RNA (Lowary and Uhlenbeck 1987), was used more recently (Ward et al. 2019) (referred to here as MS2-V2).

**Figure 2.**
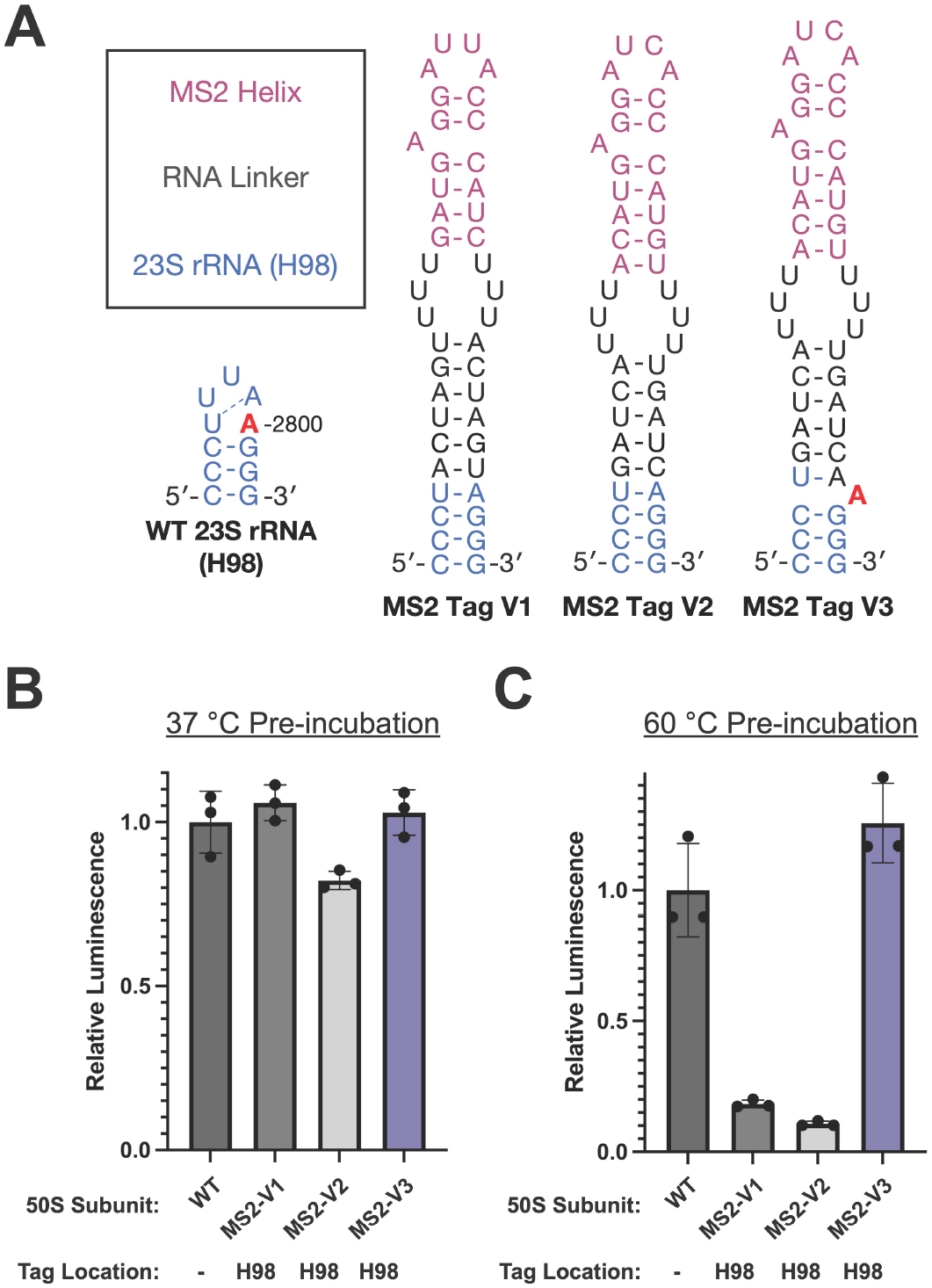
Design and activity of 50S subunits with MS2 tags inserted into H98. A) The secondary structure of H98 in WT 23S rRNA or ribosomes with MS2 tags. The sequence of MS2-V1 and MS2-V2 are from (Youngman and Green 2005) and (Ward et al. 2019) respectively. Addition of an adenosine in MS2-V3 at a position equivalent to A2800 is proposed to mimic the sequence of the WT 23S rRNA (A2800 in red). 50S subunits were preincubated at B) 37°C or C) 60°C before their addition to the nanoLuciferase *in vitro* translation assay. Data is normalized to WT subunits with no MS2 tag. All error bars are the standard deviation of three experimental replicates.

It was previously shown that 50S subunits with an MS2-V2 tag inserted into H98 are destabilized compared to WT subunits (Nissley et al. 2023). We hypothesized that the insertion of an MS2 tag into H98 disrupts the tertiary interactions between H1, H94, and H98 and causes global destabilization of the 50S subunit. In the WT ribosome structure, nucleotide A2800 in H98 interacts with bases near H1, bridging H1 and H98. Although A2800 is not removed during the insertion of MS2-V1 or V2, the extension of the helix and the loss of A2799 in the H98 apical loop could drive A2800 into a Watson-Crick base pair with U2796 in the H98 helix rather than remaining flipped out to interact with H1 and H94. To test whether we could improve the MS2 tag design, we added an additional adenosine into a position synonymous to A2800 (MS2-V3). This design would allow the additional unpaired adenosine to interact with H1 nucleotides and restore the canonical structure of this region.

### MS2-V3 restores stability to the *E. coli* LSU

To test whether the loss of A2800 in MS2 tags was the cause for destabilization of the 50S subunit, we tested the activity of 50S subunits with MS2-V1, V2, or V3 tags using an *in vitro* nanoluciferase (nLuc) translation assay. As was shown previously, the insertion of an MS2-V1 tag does not affect *in vitro* ribosome activity at 37°C (Youngman and Green 2005; Ward et al. 2019) (**Figure 2B**). We do observe a slight decrease in the activity of 50S subunits with MS2-V2 tags, however this may be due to the additional incubation step at 37°C before the *in vitro* translation assay (Methods). Notably, 50S subunits with a MS2-V3 insertion have similar levels of activity to untagged WT 50S subunits.

While ribosome activity at 37°C is not affected by the insertion of MS2 tags in H98, we also investigated the ability of 50S subunits to withstand a heat treatment step before the *in vitro* translation assays. After pre-incubation at 60°C, subunits with MS2-V1 and V2 inserted in H98 exhibit only 18% and 11%, respectively, of the activity of untagged WT subunits (**Figure 2C**). This indicates that subunits with MS2-V1 and V2 tags are thermodynamically destabilized compared to WT subunits. In contrast, 50S subunits with MS2-V3 tags demonstrate WT levels of activity after incubation at 60°C (**Figure 2C** and **Figure S1**). This suggests that loss of the A2800 interaction plays a role in the destabilization of subunits with MS2-V1 and MS2-V2 tags.

We examined the H1-H94-H98 interaction in the cryo-EM structures of a WT 70S ribosome (Watson et al. 2020) and ribosomes with an MS2-V2 tag (Nissley et al. 2023). In the WT ribosome, there is clear density for G9 (H1), A10 (H1), and U2629 (H94) sugars and bases. There is also clear density for the A2800 (H98) base despite poor density for the rest of the apical region of H98 (**Figure 3A**), indicating that while H98 is dynamic on the surface of the ribosome, A2800 is rigidly docked through contacts with H1 nucleotides. In contrast, the ribosome with MS2-V2 inserted in H98, lacks density for A2800 and displays weak density for A10 suggesting that this nucleotide is more dynamic. While A2800 is not deleted in the MS2-V2 tag, the lack of density at its canonical position suggests that the adenosine forms a Watson-Crick base pair with U2796 in the H98 helix. The RNA helical extension of H98 in the MS2-V2 tag may energetically favor the A2800-U2796 base pair and force A2800 out of its canonical flipped position observed in the WT structure.

**Figure 3.**
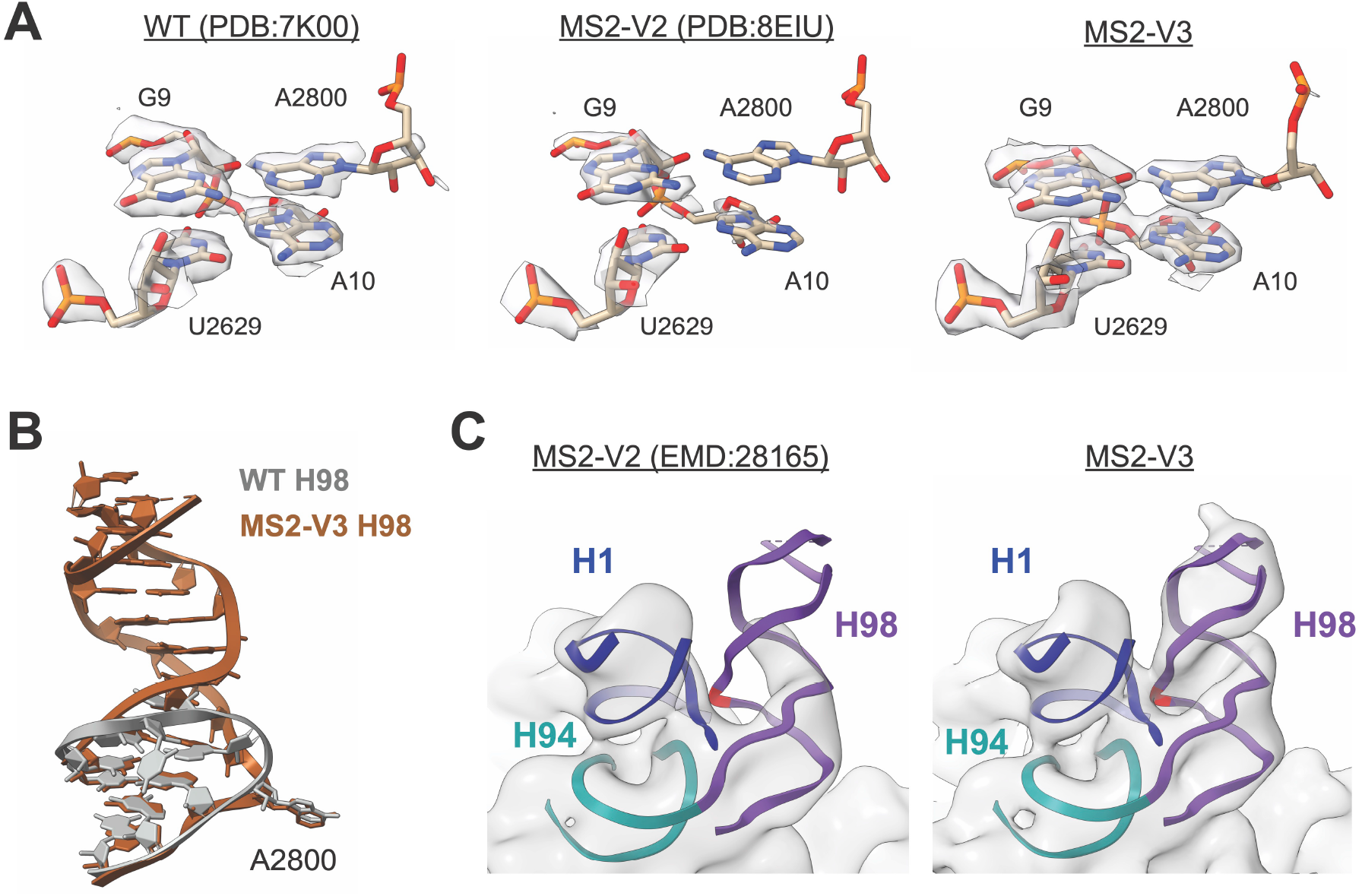
Structure of the *E. coli* 70S ribosome with MS2-V3 inserted in H98. A) Model and density for the H1-H94-H98 nucleotide quartet in the WT 70S ribosome (left, PDB:7K00 EMD:22586), 70S ribosome with MS2-V2 inserted in H98 (middle, PDB:8EIU EMD:28165), and 70S ribosome with MS2-V3 inserted in H98 (right). An additional *B*-factor of 15 Å^2^ was applied to the cryo-EM maps. B) Overview of H98 in the WT 70S ribosome (grey, PDB:7K00) and the helix upon insertion of MS2-V3 in H98 (brown). C) Low-pass filtered cryo-EM maps for MS2-V2 (EMD:28165) and MS2-V3 overlayed on the MS2-V3 model for H1, H94, and H98. Cryo-EM maps were low-pass filtered to 10 Å. Nucleotide A2833 (equivalent to A2800) is marked in red.

To determine how the addition of an adenosine residue in MS2-V3 affects the structure of the H1-H94-H98 region, we solved the structure of an MS2-V3 tagged *E. coli* 70S ribosome with fMet-tRNA^fMet^ in the P site to a global resolution of 1.8 Å resolution. This high-resolution structure enabled the visualization of individual nucleotides on the surface of the ribosome, including some of those in H1 and H98 where local resolutions range from 2.2-2.4 Å (**Figure S2**). Additionally, we utilized 3D-variability analysis in cryoSPARC to better resolve the extended MS2 tag and could model the helix into lower-resolution helical density through the poly-uridine stretch.

Although the apical MS2 helix and MS2 coat protein are likely too dynamic to be visualized, H98 with the MS2-V3 insertion adopts roughly the same conformation as WT H98, with the linker helix extending away from the surface of the 50S subunit (**Figure 3B**). Importantly, there is clear density for both A10 and the additional adenosine at a position equivalent to 2800 in the WT ribosome (2833 with the MS2-V3 tag) (**Figure 3A**). The additional adenosine is in a similar position to A2800 in the WT structure and forms similar hydrogen bonds with the other quartet nucleotides (**Figure 3B** and **Figure S3**).

To further understand how the addition of the adenosine affects the extended H98, we looked at low-pass filtered maps for ribosomes with MS2-V2 and MS2-V3 tags (**Figure 3C**). With MS2-V2 tags, there is a lack of density around canonical location of A2800 and much of the linker helix, suggesting that once H1-H98 contacts are lost, H98 becomes highly dynamic. Upon the addition of an adenosine in MS2-V3 tags, there is a recovery of density for the extended H98. Thus, in the ribosome with an MS2-V3 tag, addition of an adenosine into the MS2 tag restores the H1-H94-H98 tertiary interaction and this recovers ribosome stability.

### Nucleotide bridges between H1, H94, and H98 are required for full LSU stability

Since the interaction between H1, H94, and H98 plays a role in ribosome stability, we carefully examined the structure of this region. The nucleotide quartet is composed of two sets of base pairs which stack on one another: a type X *trans* Watson-Crick—sugar edge base pair between G9 and A2800 and a type XXIV *trans* Watson-Crick—Hoogsteen base pair between U2629 and A10. Additionally, N1 of G9 and N6 of A2800 interact with the 2′-OH of U2629 and G9, respectively (**Figure 4A**).

**Figure 4.**
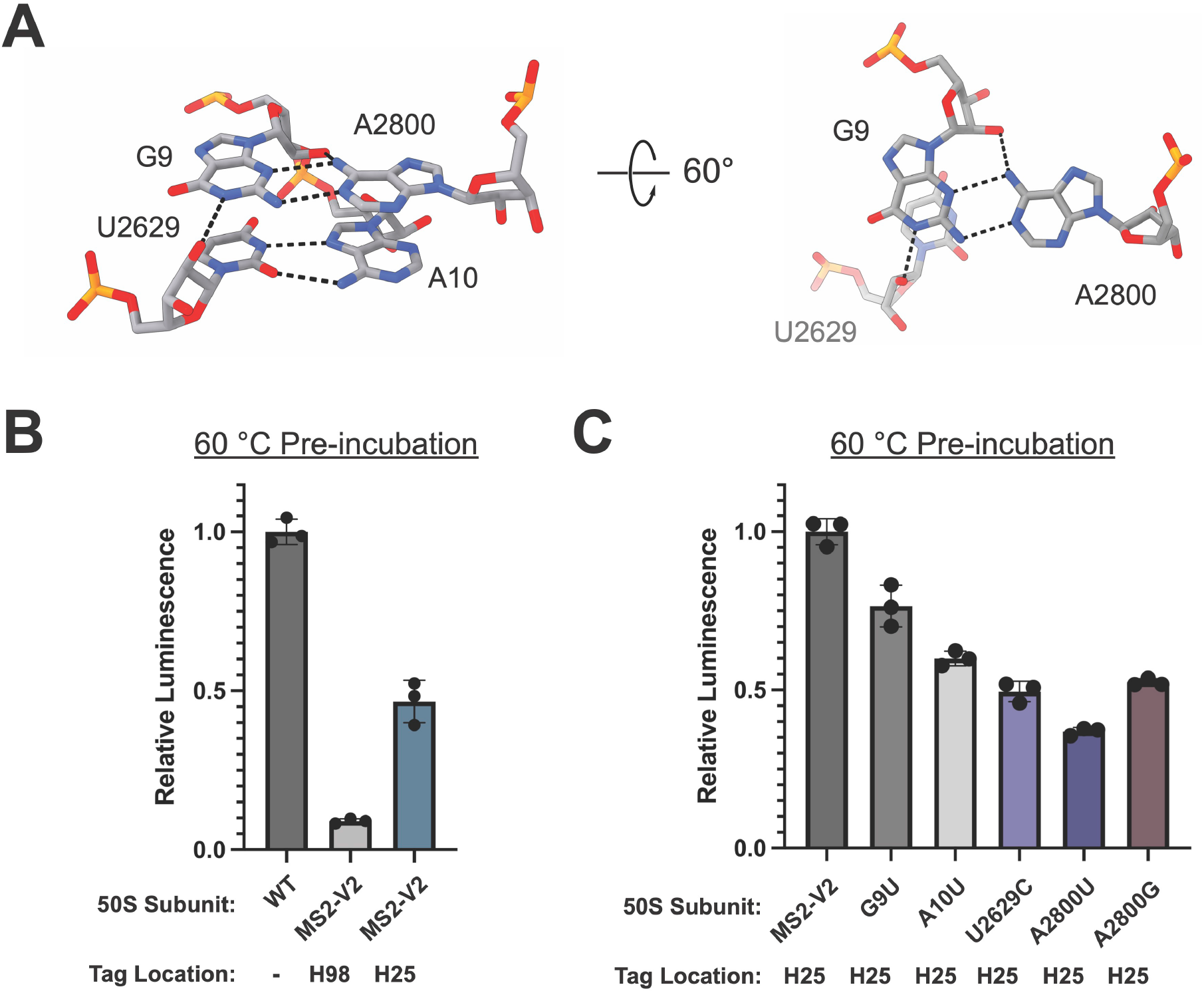
Base pairing and stacking in the H1-H94-H98 nucleotide quartet. A) Overview of the nucleotide quartet in the WT 70S ribosome. Hydrogen bonds between the bases and sugars are shown in black. B) nLuc *in vitro* translation assay for WT (no tag), MS2-V2 inserted into H98, and MS2-V2 inserted into H25 after pre-incubation of 50S subunits at 60°C. Data is normalized to untagged 50S subunits. C) nLuc *in vitro* translation assay for 50S subunits with MS2-V2 inserted into H25 and mutations in the H1-H94-H98 quartet after pre-incubation of 50S subunits at 60°C. Data is normalized to subunits with MS2-V2 inserted into H25. All error bars are the standard deviation of three experimental replicates.

To further interrogate the interactions between H1, H94, and H98, we inserted the MS2-V2 tag into H25, as a streptavidin binding aptamer was previously placed in H25 for the affinity purification of *E. coli* ribosomes (Leonov et al. 2003). By moving the MS2 tag into H25, we were able to maintain the WT sequence of H98 to determine how the native contacts in this region affect the stability of the LSU. The insertion of an MS2 tag into H25 has a slight negative effect on ribosome stability but no effect on activity at 37°C (**Figure 4B** and **Figure S4**). This result suggests that the extension of solvent accessible helices on the surface of the ribosome can have unexpected effects on ribosome folding and stability.

Utilizing the MS2 tag inserted into H25, the following 50S mutant subunits were purified: G9U, A10U, U2629C, A2800U, and A2800G. Purine to uridine mutations (G9U, A10U, and A2800U) were made to disrupt base pairing and base stacking in the H1-H94-H98 tertiary contact as uridine has higher (less favorable) base stacking free energies than purine nucleotides (Frechet et al. 1979; Hayatshahi et al. 2018). A U2629C mutation was hypothesized to make minor perturbations to the U2629-A10 pair while maintaining the smaller pyrimidine ring size.

Mutations in the quartet nucleotides did not decrease ribosome activity at 37°C but negatively affected ribosome stability (**Figure 4C** and **Figure S4**). The G9U and A10U mutant subunits had the highest stability compared to the other quartet mutations. It is likely that both nucleotides work together to fully link H1 to H94 and H98, and that the mutation of one residue does not completely abolish the interaction. Mutations to U2629 and A2800 led to the largest negative effects on ribosome stability. The mutation of these residues would prevent H94 or H98 from interacting with H1, which indicates that the loss of either of these contacts decreases ribosome stability. Subunits with an A2800U mutation had the lowest activity after pre-incubation at 60°C, suggesting that the H98-H1 interaction plays an important role in the stability of the *E. coli* LSU. When the nucleotides that base-pair (G9U) or base stack (A10U) with A2800 are mutated, there is also a loss in ribosome stability.

Additionally, a subunit with an A2800G mutation displayed slightly higher levels of activity after pre-incubation at 60°C than one with an A2800U mutation. The guanosine at position 2800 would still be able to form a purine-purine base stack with A10 but would not be able base-pair with the sugar edge of G9, indicating that base-stacking in this region likely plays an important role in nucleotide engagement.

## DISCUSSION

Here we show that the insertion of MS2 tags into H98 of 23S rRNA in the *E. coli* LSU using an uninterrupted RNA double helical extension (Youngman and Green 2005; Ward et al. 2019) disrupts contacts between H1, H94, and H98 and destabilizes the LSU. Stability is recovered upon the addition of an adenosine into H98 MS2 tags at a position equivalent to nucleotide 2800 in 23S rRNA of the WT subunit, which restores the H1-H94-H98 interaction. It has been shown that affinity handles inserted into helices on the surface of the *E. coli* LSU can have negative effects on ribosome activity, assembly, and cellular growth (Leonov et al. 2003; Hesslein 2004). The affinity tag described here builds on an MS2 tag design that does not affect cellular growth or ribosome activity (Youngman and Green 2005) and now also achieves WT levels of thermodynamic stability. Additionally, the design principles outlined here can be applied to the placement of affinity tags in H98 of other bacterial LSUs, where maintenance of H1-H94-H98 interactions should be prioritized. Maintaining stability in the LSU could be important when studying ribosome mutations elsewhere that affect assembly, RNA folding, and global stability, since conferring additional instability from the MS2 affinity tag could lead to inactive ribosomes.

H98 is a poorly conserved helix in 23S rRNA that is absent in ribosomes from many organisms (Matadeen et al. 2001). Since H1, H94, and H98 play a role in the stability of the *E. coli* LSU, we wondered whether these helices play similar roles in other bacterial ribosomes. We examined the region surrounding H1 in various high resolution structures of bacterial ribosomes that contain H98 (**Table 1**) (Basu et al. 2022; Morgan et al. 2022; Syroegin et al. 2023; Kaminishi et al. 2015; Crowe-McAuliffe et al. 2021b, 2021a; Halfon et al. 2019; Murphy et al. 2020; Cui et al. 2022; Mishra et al. 2018). Out of the surveyed structures, the ribosomes from two organisms, *P. aeruginosa* and *E. faecalis*, have interactions between H1, H94, and H98 that form the same structure as in the *E. coli* ribosome. However, other organisms possess diverse sets of contacts between these helices that differ from the ones found in *E. coli* (**Figure 5**). For example, the interactions between the helices are more expansive in *M. tuberculosis* and *T. thermophilus* than in *E. coli. T. thermophilus* ribosomes utilize complex hydrogen bond networks and base stacking that involves both the 5′ and 3′ strands of H1, to bridge H1 with H94 and H98 (**Figure 5**).

**Table 1.** Interactions between rRNA helices H1, H94, and H98 from bacterial ribosomes with known structures. The identities of RNA nucleotides from these helices which interact are listed. H1 nucleotides on the 3′ strand of 23S rRNA are shown in parenthesis.

| Organism | Phylum | H1 Interacting Nucleotides | H94 Interacting Nucleotides | H98 Interacting Nucleotides |
| --- | --- | --- | --- | --- |
| *Escherichia coli* | Pseudomonadota | G A | U | A |
| *Pseudomonas aeruginosa* | Pseudomonadota | G A | U | A |
| *Actinetobacter baumannii* | Pseudomonadota | A A | U | U |
| *Thermus thermophilus* | Deinococcota | U G (U) | A | A |
| *Deinococcus radiodurans* | Deinococcota | U A | A | - |
| *Bacillus subtilis* | Bacillota | U A | A | - |
| *Listeria monocytogenes* | Bacillota | U A | A | - |
| *Staphylococcus aureus* | Bacillota | U A | A | - |
| *Enterococcus faecalis* | Bacillota | G A | U | A |
| *Mycobacterium tuberculosis* | Actinomycetota | G U (A) | C | G |
| *Mycolicibacterium smegmatis* | Actinomycetota | U U | C | - |

**Figure 5.**
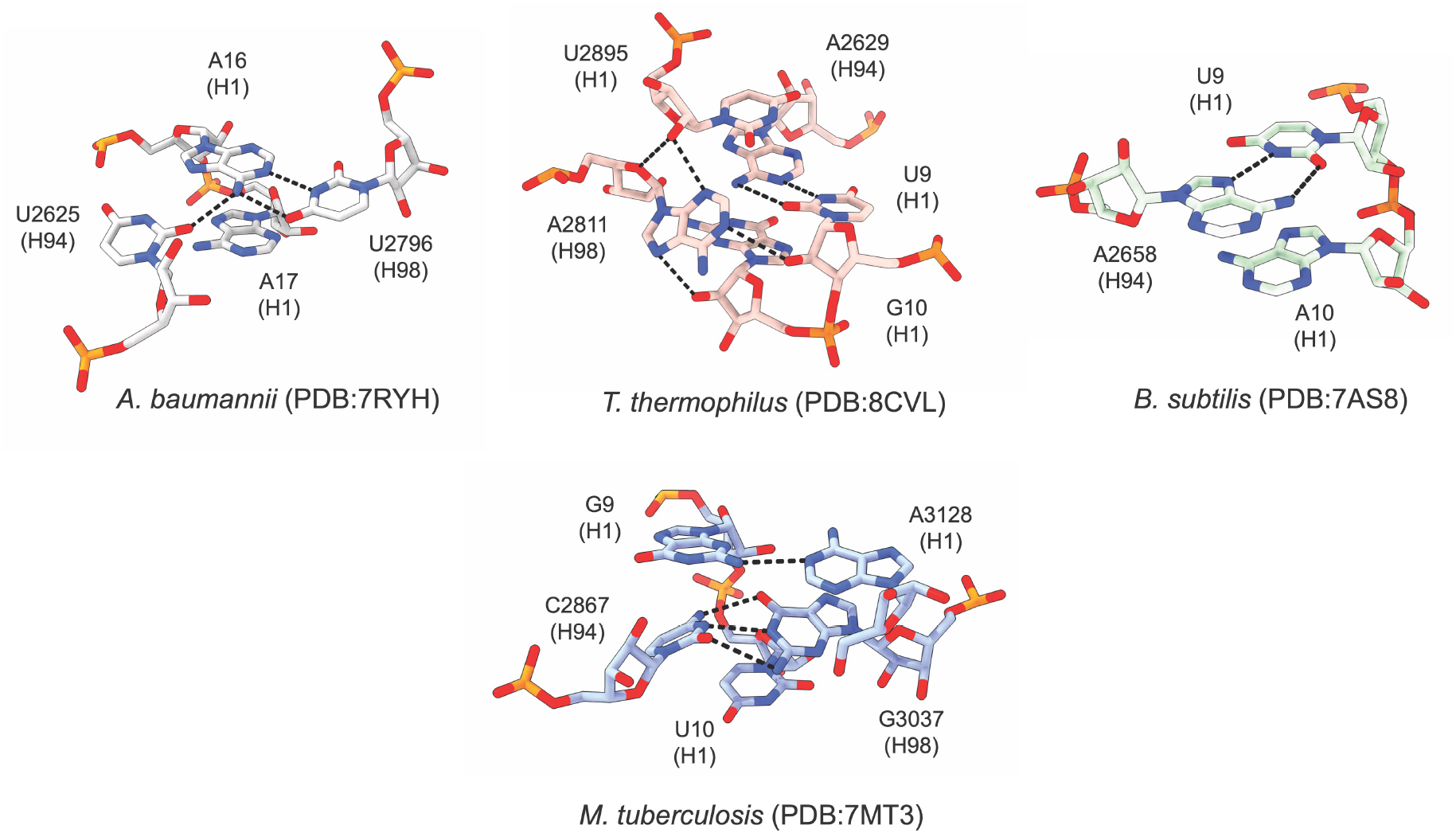
Interactions between H1, H94, and H98 in various bacterial ribosomes. Nucleotides from H1, H94, and H98 that interact with one another are displayed. Hydrogen bonds between the bases and sugars are shown in black.

This is suggestive of a stronger interaction that could be advantageous at the higher growth temperatures of *T. thermophilus*. In all the surveyed ribosome structures there are interactions between the 5′ strand of H1 and a variable nucleotide at the base of H94.

However, H98 does not interact with H1 in many of the structures, such as in *B subtilis*. In these bacteria, the H1-H94 interaction or other compensating interactions may be sufficient for ribosome stability.

Previously it was shown that some bacteria lose H1 during the maturation of the LSU and that the loss of H1 is correlated with the evolutionary loss of H98 (Shatoff et al. 2021).

This finding hinted at an unknown synergistic role between the two helices and that H1 became redundant after the evolutionary loss of H98. It was hypothesized that H98 may play a role in protecting H1 from RNase cleavage (Shatoff et al. 2021), and our data suggests that H98 plays an additional role thermodynamically stabilizing the ribosome through interactions with the 5′ strand of H1. As the ribosomes of many organisms lack H98 and some bacteria cleave H1 during LSU assembly, there may be compensating interactions that replace the H1-H94-H98 interaction in these ribosomes. For example, in the *Flavobacterium johnsoniae* ribosome, which lacks H1 in the mature LSU, the 5′ and 3′ ends of the 23S rRNA interact with L22 and L32, respectively (Jha et al. 2021). L22 and L32 form a protein-protein interaction in the *F. johnsonjae* ribosome, which may play a synonymous role to the H1-H94-H98 RNA-RNA interaction in *E. coli* as it bridges the 5′ and 3′ ends of 23S rRNA.

RNA intrinsically forms compact structures and it has been shown that the 5′ and 3′ ends of long RNAs are in close proximity (Lai et al. 2018; Leija-Martínez et al. 2014; Yoffe et al. 2011), likely playing a role in RNA folding and stability. This observation has also been made for the termini of many proteins, and in certain proteins the loss of the N-terminal—C-terminal interaction leads to protein destabilization or unfolding (Krishna and Englander 2005). RNAs with compact structures, such as rRNA and other ribozymes, have closer end to end distances than RNAs with less stable structures (Vicens et al. 2018). While H1 bridges the 5′ and 3′ ends of 23S rRNA, additional interactions between the 5′ end of H1 and terminal helices H94 and H98 on the 3′ end likely work to further tether the ends of the 23S rRNA. The interactions between helices on the ends of 23S rRNA likely stabilize the fold of certain bacterial ribosomal large subunits and may more broadly represent a method for stabilizing the fold of highly structured RNAs.

## MATERIAL AND METHODS

### Cloning and plasmid design

A modified pLK35 plasmid (Douthwaite et al. 1989), containing a tac promoter followed by the rrnB operon, which encodes the MS2-tagged 23S rRNA, was used for mutagenesis. Point mutations, insertions, and deletions were made using either the Q5 mutagenesis kit (NEB) or the InFusion cloning kit (Takara Bio) and the corresponding primer sets (**Table S1**). MS2 tags inserted in H98 were placed between 23S rRNA bases 2796 and 2800, with the concomitant removal of nucleotides 2797-2799. To insert an MS2 tag into H25, the MS2-V2 sequence was placed between 23S rRNA bases 544 and 549, with the removal of nucleotides 545-548. The following DNA sequences were inserted for each tag design:

MS2-V1: 5′-ACTAGTTTTGATGAGGATTACCCATCTTTACTAGT-3′

MS2-V2: 5′-GATCATTTACATGAGGATCACCCATGTTTTTGATC-3′

MS2-V3: 5′-GATCATTTACATGAGGATCACCCATGTTTTTGATCA-3′

### Ribosome expression and purification

Mutant *E. coli* 50S ribosomal subunits were expressed and purified as previously described (Nissley et al. 2023). Briefly, *E. coli* NEB Express I^q^ cells were transformed with the corresponding pLK35 plasmid. Transformants were grown in 3 L of LB broth containing 100 μg/mL ampicillin at 37°C. Cultures were induced with 0.5 mM isopropyl β-D-1-thiogalactopyranoside once they reached OD_600_=0.6 and were incubated for an additional 3 hours. Cells were pelleted, resuspended in 50 mL buffer A (20 mM Tris-HCl pH 7.5, 100 mM NH_4_Cl, 10 mM MgCl_2_, 0.5 mM EDTA, 2 mM dithiothreitol (DTT)) with Pierce protease inhibitor (ThermoFisher), and lysed by sonication. The lysate was clarified by centrifugation and then loaded onto a sucrose cushion containing 24 mL of buffer B (20 mM Tris-HCl pH 7.5, 500 mM NH_4_Cl, 10 mM MgCl_2_, 0.5 mM EDTA, 2 mM DTT) with 0.5 M sucrose and 17 mL of buffer C (20 mM Tris-HCl pH 7.5, 60 mM NH_4_Cl, 6 mM MgCl_2_, 0.5 mM EDTA, 2 mM DTT) with 0.7 M sucrose in Type 45 Ti tubes (Beckman-Coulter). Ribosomes were pelleted by centrifugation at 27,000 rpm (57,000 xg) for 16 hours at 4°C. The pellets were then resuspended in dissociation buffer (buffer C with 1 mM MgCl_2_), which dissociates 70S ribosomes into 30S and 50S subunits.

MS2-tagged 50S subunits were then purified using a maltose binding protein (MBP)-MS2 fusion protein, which was purified as previously described (Ward et al. 2019). A 5 mL MBP Trap column (Cytiva) at 4°C was washed with 5 column volumes (CV) of MS2-150 buffer (20 mM HEPES pH 7.5, 150 mM KCl, 1 mM EDTA, 2 mM 2-mercaptoethanol) and then 10 mg of MBP-MS2 protein was loaded slowly onto the column. The column was then washed with 5 CV of buffer A-1 (buffer A with 1 mM MgCl_2_), and crude ribosomes were loaded onto the column. The column was washed with 5 CV of buffer A-1, 10 CV of buffer A-250 (buffer A with 250 mM NH_4_Cl and 1 mM MgCl_2_), and then tagged ribosomal subunits were eluted with a 10 CV gradient of 0-10 mM maltose in buffer A-1. The purified 50S subunits were concentrated in 100 kDa cut off spin filters (Millipore), quantified with an approximation of 1 A_260_= 36 nM, and flash frozen in liquid nitrogen. WT untagged 30S and 50S ribosomal subunits were purified as previously described (Nissley et al. 2023).

### Mutant ribosome purity assay

To determine the levels of WT untagged 50S subunit contamination after mutant ribosome purification, semiquantitative PCR was utilized as previously described (Ward et al. 2019) with adaptations. Roughly 50 pmol of purified mutant ribosomes were denatured at 95°C for 3 minutes. LiCl was added to a final concentration of 5 M to precipitate the rRNA, which was subsequently resuspended in RNase-free water. Primers MS2_H98_quant_R or MS2_H25_quant_R, for mutant ribosomes with MS2 tags in either H98 or H25 respectively, were used to reverse transcribe a region of 23S rRNA using AMV reverse transcriptase (Promega). The region of cDNA containing the MS2 tag was amplified via PCR using primer pairs MS2_H98_quant_F and MS2_H98_quant_R or MS2_H25_quant_F and MS2_H25_quant_R. PCR products were resolved on a 10% polyacrylamide-TBE gel (Invitrogen) and visualized with SYBR gold stain (ThermoFisher). DNA bands were quantified with ImageJ software (Schneider et al. 2012).

### *In vitro* nanoluciferase translation endpoint assay

50S subunits were diluted to 1.4 μM in buffer A with a final concentration of 10 mM MgCl_2_. The diluted subunits were incubated for 15 minutes at the indicated temperature and then incubated for an additional 15 minutes at room temperature to cool. An *in vitro* translation reaction was then assembled using the PURExpress system (NEB) with the following: 3.2 μL solution A (NEB), 1 μL factor mix (NEB), 250 nM pre-incubated 50S subunit, 500 nM WT 30S subunit, 1 U/μL RNase inhibitor (NEB), and 10 ng/μL of a plasmid containing a T7 promoter followed by the nanoluciferase (nLuc) gene (final volume of 8 μL). The reaction was incubated for 1 hour at 37°C and then 2 μL of the reaction was mixed with 30 μL of nLuc buffer (20 mM HEPES pH 7.5, 50 mM KCl, and 10% glycerol) and a 1:50 dilution of Nano-Glo substrate (Promega). All 32 μL were then placed in a 384 well plate and luminescence was measured in a Spark Plate Reader (Tecan). Experimental triplicates were measured and averaged for each *in vitro* translation reaction.

### MetRS and MTF expression and purification

Plasmids containing a T7 promoter followed by the gene for *E. coli* methionyl-tRNA synthetase (MetRS) or methionyl-tRNA formyltransferase (MTF) and a 6x-His tag were transformed into *E. coli* BL21 (DE3) Codon+ RIL cells. Overnight cultures were diluted in ZYM-5052 autoinducing media (Studier, F. W. 2014) and grown overnight at 37°C. Cells were then pelleted by centrifugation, resuspended in lysis buffer (20 mM Tris pH 7.8, 150 mM NaCl, 5 mM imidazole, 0.5 mM EDTA), and lysed by sonication. The lysate was clarified by centrifugation at 25,000 xg (JA-20 rotor, Beckman) at 4°C for 30 minutes. The supernatant was applied to a 5 mL HisTrap column (Cytiva) at 4°C and the column was washed with 5 CV of lysis buffer with 23 mM imidazole. Protein was eluted from the column using a linear gradient of 20 CV lysis buffer from 23-500 mM imidazole. The eluted protein was then dialyzed against 50 mM HEPES pH 7.5, 100 mM KCl, 10 mM MgCl_2_, 7 mM β-mercaptoethanol (BME), and 30% glycerol, concentrated, and stored at -80°C.

### fMet-tRNA^fMet^ preparation

tRNA^fMet^ with a C1G mutation and a modified terminal 3′-NH_2_-ATP was *in vitro* transcribed and modified as previously described (Nissley et al. 2023). fMet-tRNA^fMet^ was prepared enzymatically as described (Walker and Fredrick 2008). Briefly, a charging reaction was prepared with the following: 10 μM NH_2_-tRNA^fMet^, 10 mM methionine, 300 μM N10-formyl-tetrahydrofolate, 10 mM ATP, 1 U/μL RNase inhibitor (NEB), 1 μM (MetRS), and 1 μM MTF in AA buffer (50 mM HEPES pH 7.5, 10 mM KCl, 20 mM MgCl_2_, and 2 mM DTT). This reaction was incubated at 37°C for 30 minutes. The tRNA was then phenol-chloroform extracted, ethanol precipitated, and resuspended in water.

### Cryo-EM sample preparation

40 pmol of MS2-V3 50S subunit and 80 pmol WT untagged 30S subunit were incubated at 37°C in buffer C with a total of 10 mM MgCl_2_ for 45 minutes. This mixture was then split and loaded on to four 15-40% (w/v) sucrose gradients in buffer D (20 mM HEPES pH 7.5, 100 mM KCl, 10 mM Mg(OAc)_2_). Gradients were centrifuged at 97,000 xg for 16 hours in a SW-41 rotor (Beckman-Coulter). An ISCO fractionation system was used to isolate the 70S fraction from each gradient (**Figure S5**). 70S factions were then combined, washed in buffer D, and concentrated in a 100 kDa cut off spin filter (Millipore).

100 nM 70S ribosome, 1 μM fMet-tRNA^fMet^, and 5 μM mRNA with the sequence 5′-GUAUAAGGAGGUAAA<u>AUG</u>UUCUAACUA-3′ (IDT), were combined in Buffer D with 15 mM MgOAc and incubated at 37°C for 45 minutes. The fMet codon is underlined in the mRNA sequence. 300 mesh R1.2/R1.3 UltraAu foil grids with a deposited layer of amorphous carbon were glow discharged in a PELCO easiGlow for 12 seconds under 0.37 mBar vacuum and with 25 mA current. 4 μL of the complex was then added to the grid, incubated for 1 minute, and then the grid was touched to three successive 100 μL drops of buffer D. Grids were plunge-frozen in liquid ethane using a Vitrobot Mark IV at 4°C with 100% humidity and the settings blot force 4 and blot time 2 s.

### Cryo-EM data acquisition, image processing, and modeling

Data was acquired as previously described (Watson et al. 2020) with adaptations (**Table S2**). Briefly, movies were collected on a 300 kV Titan Krios microscope with a BIO-energy filter and Gatan K3 camera. The super-resolution pixel size was set to 0.405 (physical pixel size of 0.81) and SerialEM (Schorb et al. 2019) was used to automate data collection. Movies were collected over a defocus range of -0.5 to -1.5 μm with an electron dose of 40 e^-^/Å^2^ split over 40 frames. Image shift was used to collect movies in a 3×3 grid of holes with two movies collected per hole.

Raw movies were Fourier-cropped to the physical calibrated pixel size (0.8248 Å) and patch motion corrected in cryoSPARC 3 (Punjani et al. 2017). CTFFind4 (Rohou and Grigorieff 2015) was used to estimate the CTFs of micrographs and ones with poor estimated CTF fit were manually excluded. Micrographs were then split into exposure groups based on the 3×3 groups during image-shift collection. Particles were picked using the cryoSPARC template picker with 70S ribosome 2D templates generated in cryoSPARC. Particles were then extracted and Fourier-cropped to 1/8 of the box size and 2D classification was performed in cryoSPARC using 100 classes. Junk particles were rejected and a second round of 2D classification was performed to further remove the remaining junk particles. Selected particles were re-extracted and Fourier-cropped to 1/4 of the box size and 3D classification was performed with heterogeneous refinement in cryoSPARC using a 70S ribosome map from (Nissley et al. 2023). Classes containing features consistent with 70S ribosomes were combined and subjected to another round of 3D classification. Classes from the second round of 3D classification that refined to high resolution were combined and particles from these classes were extracted at the full box size. The particles were then subjected to homogeneous refinement in cryoSPARC with per-particle defocus optimization, per-group CTF parameter optimization, and Ewald sphere correction. Focus refinements were run separately on the 50S and 30S subunits with local refinement in cryoSPARC (**Figure S6**). For modeling, a composite map was constructed by combining the maps from the 50S and 30S focused refinements. PDB 7K00 (Watson et al. 2020), which was used as an initial 70S ribosome model, was aligned to the composite map in ChimeraX (Pettersen et al. 2021). Real space refinement in PHENIX (Liebschner et al. 2019) was used to refine the model, and the manual adjustments were made to the model as needed in COOT (Casañal et al. 2020) (**Table S3**). To better model the MS2 tag, particles were Fourier-cropped to 1/2 the box size and the region containing H1 and H98 was masked and subjected to 3D variability analysis in cryoSPARC (**Figure S6**). Each of the volume series were inspected and the map which contained the best density for the MS2 tag and showed characteristics of an RNA helix was selected. The MS2 tag was modeled into this map using secondary structure constraints in PHENIX.

## DATA DEPOSITION

Atomic coordinates have been deposited with the Protein Data Bank under accession code 8FTO. Cryo-EM maps have been deposited with the Electron Microscopy Data Bank under the accession codes EMD-29449 (composite map), EMD-29483 (70S global map), EMD-29484 (50S focused refinement map), and EMD-29485 (30S focused refinement map).

## Supporting information

PDB coordinates

cryo-EM map around H1-H94-H98 region

## SUPPLEMENTARY MATERIAL

Supplementary material is available for this article.

## ACKNOWLEDGEMENTS

We thank Dan Toso and Paul Tobias for help with cryo-EM data collection, Chandrima Majumdar for providing tRNA for cryo-EM sample preparation, and Fred Ward for help with protein purification. This work was funded by the NSF Center for Genetically Encoded Materials (C-GEM), CHE-2002182.

