## Supplementary Figures and Tables for "Interactions between terminal ribosomal RNA helices stabilize the *Escherichia coli* large ribosomal subunit"

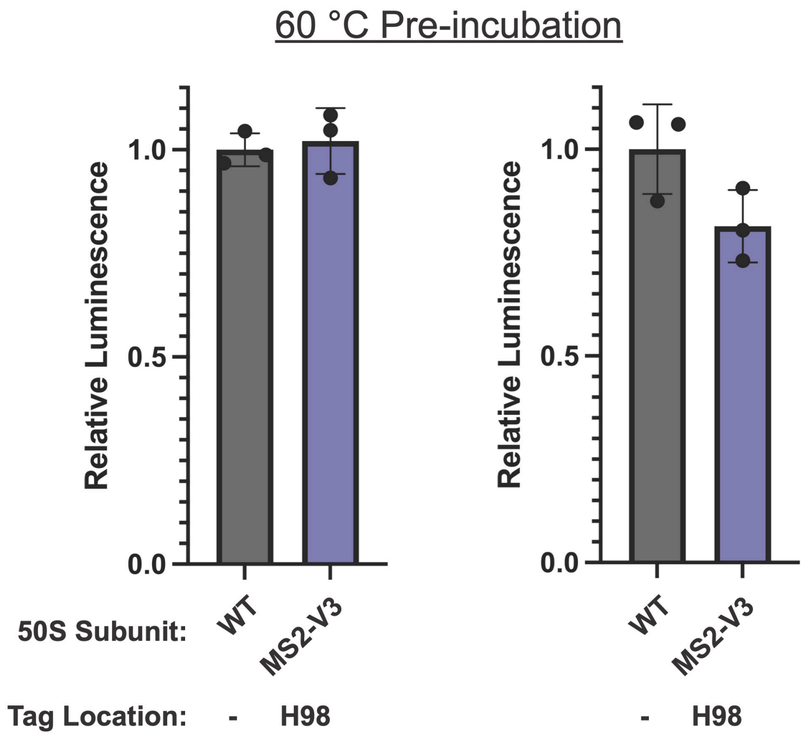


**Figure S1: Additional replicates of nanoluciferase *in vitro* translation assays for 50S subunits with MS2-V3 tags inserted into H98.** 50S subunits were preincubated at 60°C before adding to the *in vitro* translation assay. Each graph represents an independent replicate and data is normalized to WT subunits with no MS2 tag. Error bars are represented as the standard deviation of three experimental replicates.

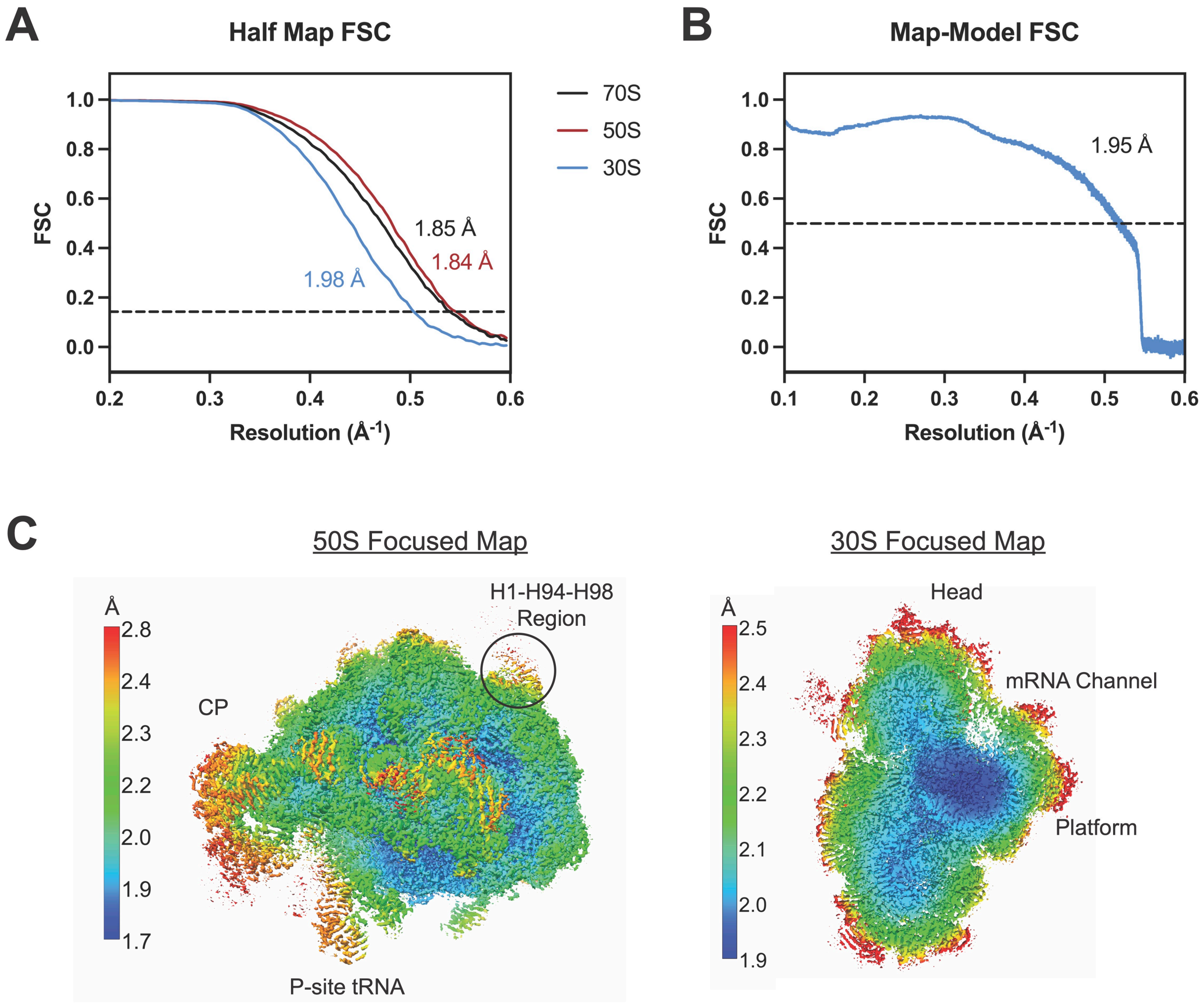


**Figure S2: Resolutions of cryo-EM maps.** A) The half map FSC resolutions (0.143 cut-off) for the global map (70S), 50S focused refinement map, and 30S focused refinement map are shown. B) The global 70S ribosome map to model FSC resolution (0.5 cut-off) is 1.95 Å. C) Local resolution is plotted on the 50S (left) or 30S (right) focused refinement maps.

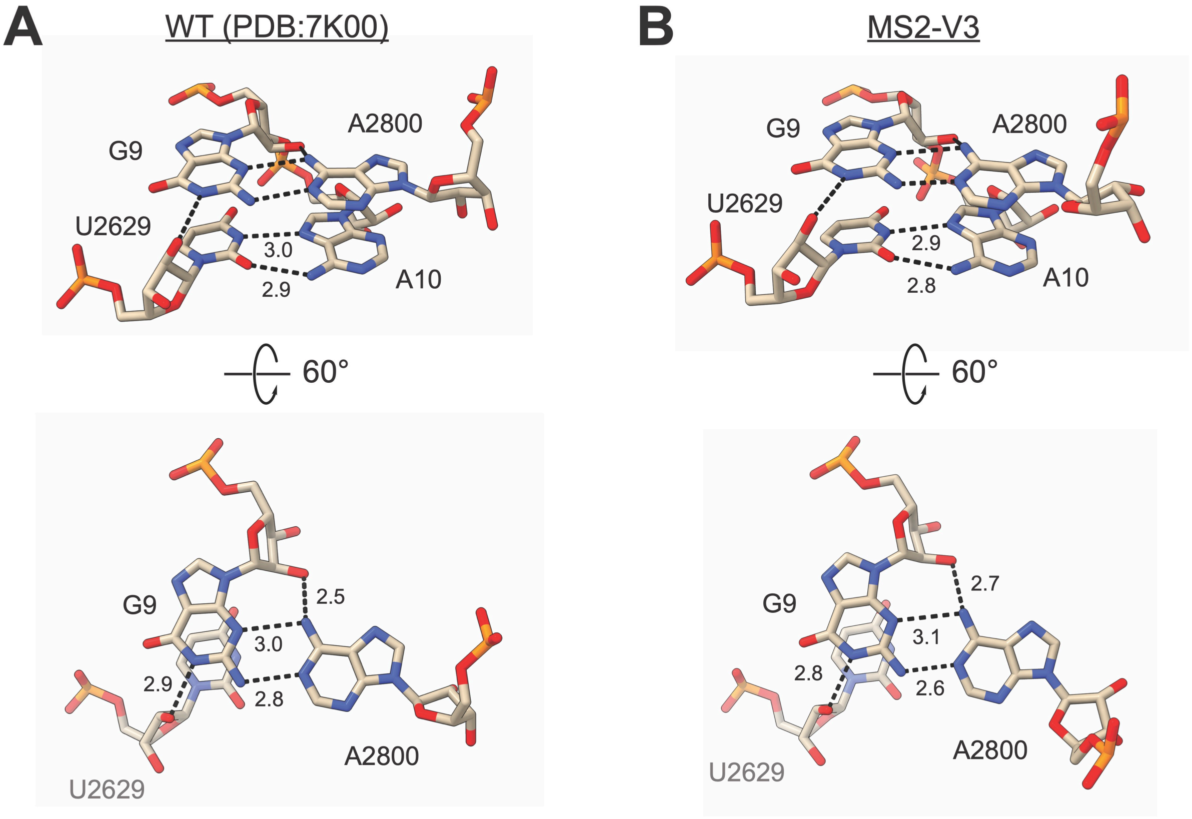


**Figure S3: Hydrogen bonding between H1, H94, and H98 nucleotides.** The nucleotides in A) WT (PDB:7K00) and B) MS2-V3 tagged ribosomes are shown, with hydrogen bonds indicated by dashed lines and distances given in Å.

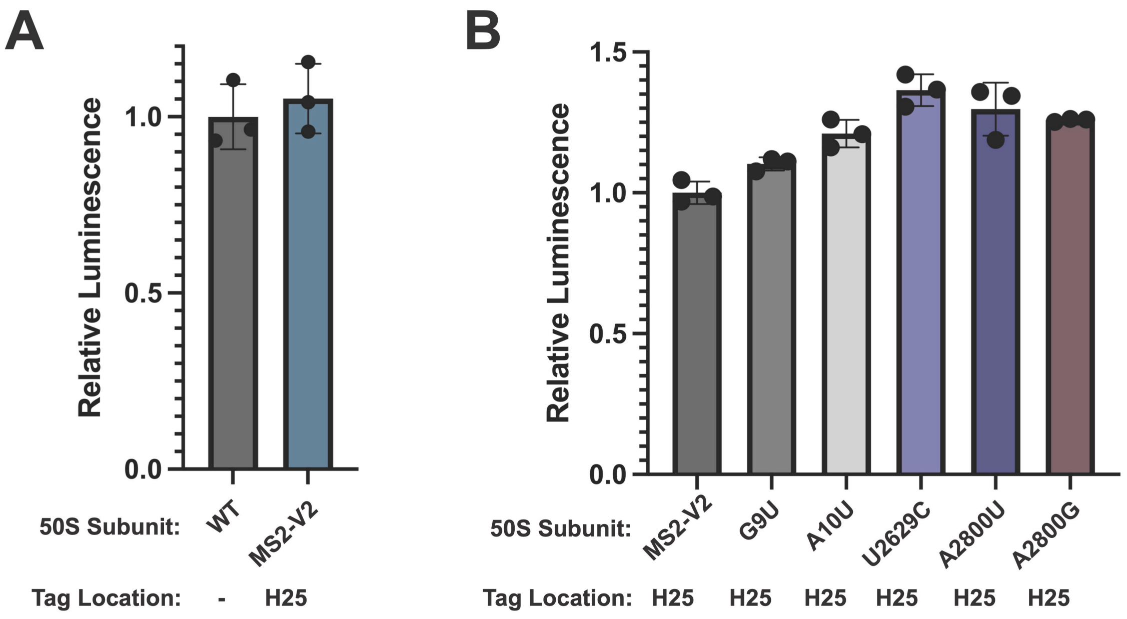


**Figure S4: Nanoluciferase *in vitro* translation assay for 50S subunits with MS2 tags with mutations in H1, H94, and H98.** A) nLuc *in vitro* translation assay for WT (no tag), and MS2-V2 inserted into H25. Data is normalized to untagged 50S subunits. B) nLuc *in vitro* translation assay for 50S subunits with MS2-V2 inserted into H25 and mutations in the H1-H94-H98 nucleotide quartet. Data is normalized to subunits with MS2-V2 inserted into H25. In these experiments, 50S subunits were not pre-incubated prior to the *in vitro* translation assay. All error bars are the standard deviation of three experimental replicates.

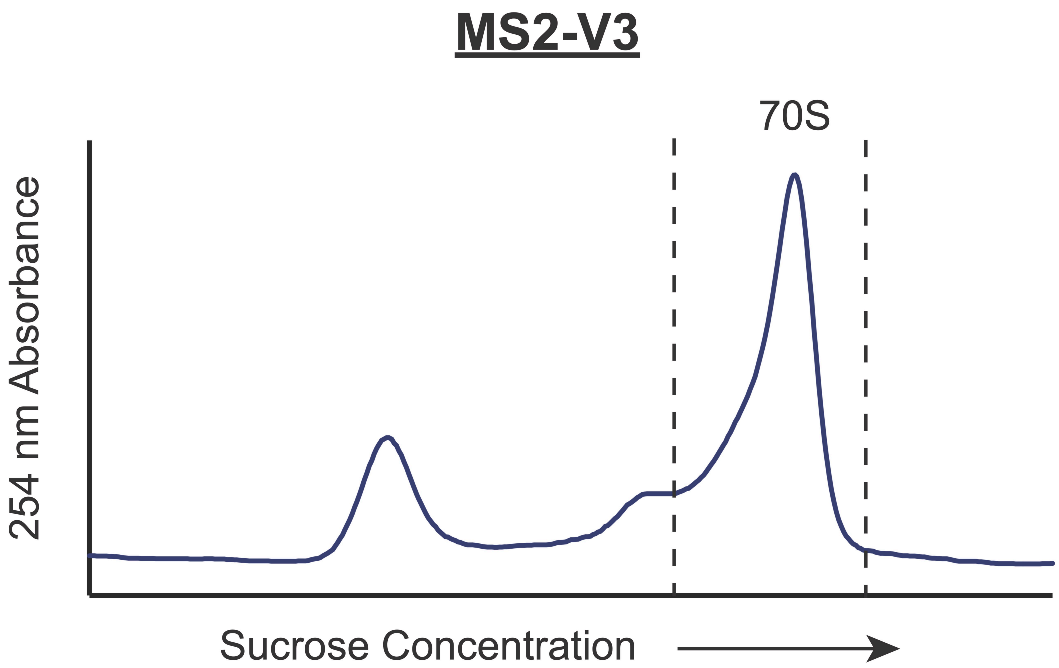


**Figure S5: Association of 50S subunits with MS2-V3 tags inserted in H98 and untagged WT *E. coli* 30S subunits.** 30S subunits (20 pmol) and 50S subunits (10 pmol) were incubated with 10 mM $MgCl_2$ and resolved on a 15-40% sucrose gradient. 70S fractions were collected between the dashed lines for cryo-EM analysis.

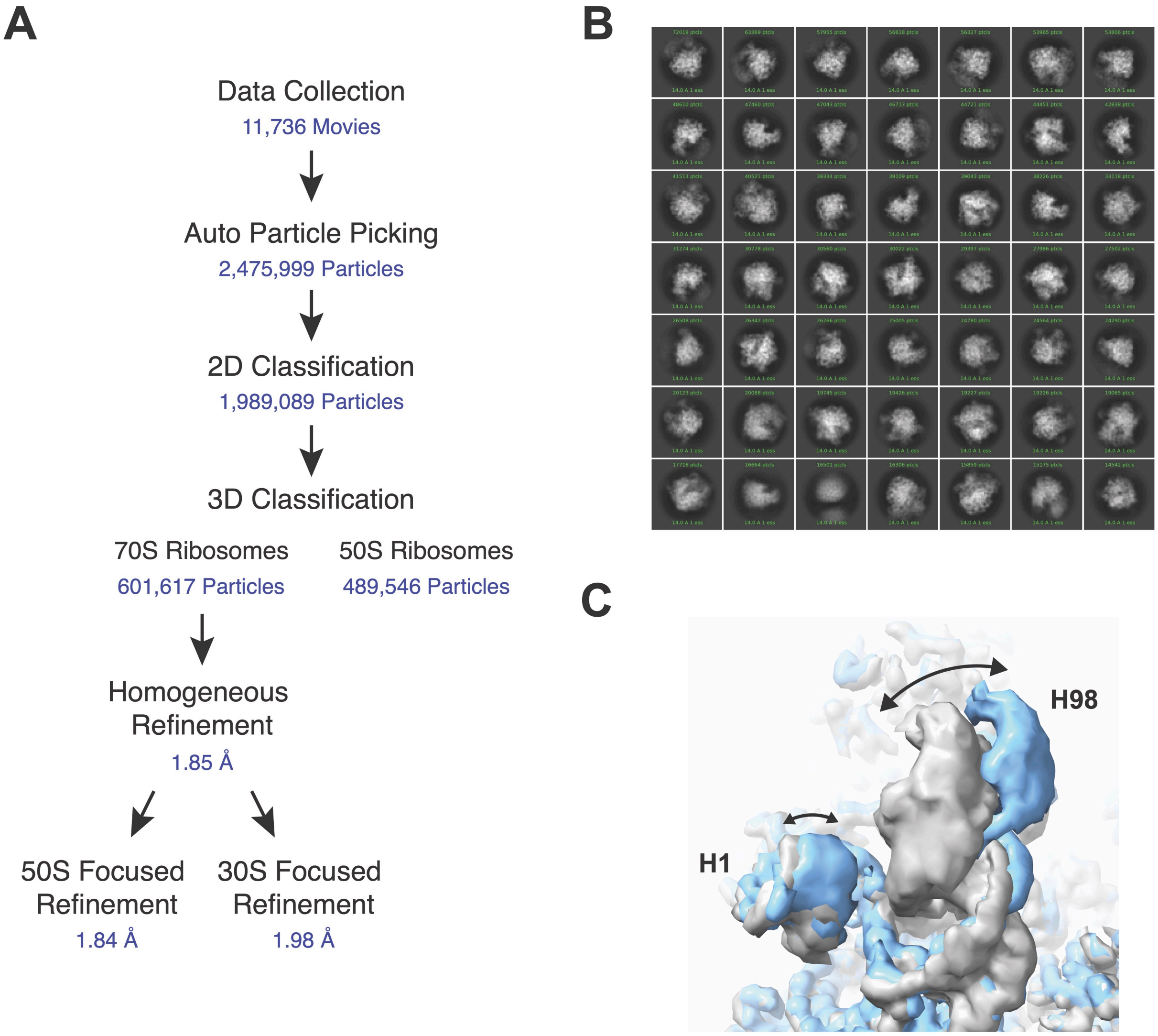


**Figure S6: Cryo-EM data processing.** A) The cryo-EM processing workflow is shown with the number of particles or half-map FSC resolution (0.143 cut-off) at each stage. B) Representative 2D classes. C) First and last frame from a component of 3D variability analysis of the region surrounding H98. Arrows demonstrate movement of H1 and H98.

**Table S1. DNA primers used in this study**

(m denotes 2'-O-Me; lowercases in the sequence indicate sites where mutations are introduced)

| **Primer Name** | **Sequence** | **Description** |
| --- | --- | --- |
| Add_A2800_H98MS2_F | TTTTGATCAAGGGTCCTGAAGGAACGTTGA | Forward primer for the addition of A2800 into MS2V2 in H98 (to make MS2V3). InFusion Kit |
| Add_A2800_H98MS2_R | GACCCTTGATCAAAAACATGGGTGATCCTCA | Reverse primer for the addition of A2800 into MS2V2 in H98 (to make MS2V3). InFusion Kit |
| Delete_MS2_H98_F | TGACCCTTTAAGGGTCCTGAAGGAACGTTGA | Forward primer for the deletion of MS2V2 from H98 (restores WT 23S rRNA). InFusion Kit |
| Delete_MS2_H98_R | ACCCTTAAAGGGTCAGGGAGAACTCATCTCG | Reverse primer for the deletion of MS2V2 from H98 (restores WT 23S rRNA). InFusion Kit |
| Insert_MS2_H25_F | cacccatgtttttgatcGCGTGTGACTGCGTACCTTTTG | Forward primer for the insertion of MS2V2 into H25. Q5 Kit |
| Insert_MS2_H25_R | atcctcatgtaaatgatcGCGTGCTCCCACTGCTTG | Reverse primer for the insertion of MS2V2 into H25. Q5 Kit |
| Insert _H98MS2_V1_F | tacccatctttactagtAGGGTCCTGAAGGAACGT | Forward primer for the insertion of MS2V1 into H98. Q5 Kit |
| Insert_H98MS2_V1_R | atcctcatcaaaactagtAGGGTCAGGGAGAACTCAT | Reverse primer for the insertion of MS2V1 into H98. Q5 Kit |
| G9U_F | GAGGTTAAGCtACTAAGCGTAC | Forward primer for G9U 23S rRNA mutagenesis (H25MS2). Q5 Kit |
| G9U_R | ACAACCCGAAGATGTTTC | Reverse primer for G9U 23S rRNA mutagenesis (H25MS2). Q5 Kit |
| A10U_F | AGGTTAAGCGtCTAAGCGTAC | Forward primer for A10U 23S rRNA mutagenesis (H25MS2). Q5 Kit |
| A10U_R | CACAACCCGAAGATGTTTC | Reverse primer for A10U 23S rRNA mutagenesis (H25MS2). Q5 Kit |

| | | |
|---|---|---|
| U2629C_F | CCGTGGGCGCcGGAGAACTGA | Forward primer for U2629C 23S rRNA mutagenesis (H25MS2). Q5 Kit |
| U2629C_R | CAGATAGGGACCGAACTGTCTC | Reverse primer for U2629C 23S rRNA mutagenesis (H25MS2). Q5 Kit |
| A2800U_F | TGACCCTTTAtGGGTCCTGAAG | Forward primer for A2800U 23S rRNA mutagenesis (H25MS2). Q5 Kit |
| A2800U_R | GGGAGAACTCATCTCGGG | Reverse primer for A2800U 23S rRNA mutagenesis (H25MS2). Q5 Kit |
| A2800G_F | TGACCCTTTAgGGGTCCTGAAG | Forward primer for A2800G 23S rRNA mutagenesis (H25MS2). Q5 Kit |
| A2800G_R | GGGAGAACTCATCTCGGG | Reverse primer for A2800G 23S rRNA mutagenesis (H25MS2). Q5 Kit |
| MS2_H98_quant_F | CTTGCCCCGAGATGAGTTCTCCC | RT-PCR primer for the ribosome purity assay with H98 MS2 tags |
| MS2_H98_quant_R | GTACCGGTTAGCTCAACGCATCGCT | Primer for amplification of cDNA in the ribosome purity assay with H98 MS2 tags |
| MS2_H25_quant_F | ACCGTGTACGTACAAGCAGTGGG | RT-PCR primer for the ribosome purity assay with H25 MS2 tags |
| MS2_H25_quant_R | CCCTTCGGCTCCCCTATTCGGTTAAC | Primer for amplification of cDNA in the ribosome purity assay with H25 MS2 tags |

**Table S2. Cryo-EM Data Collection and Processing**

| | |
|---|---|
| Magnification | 105,000 |
| Voltage (kV) | 300 |
| Electron Exposure ($e^-$/Å$^2$) | 40 |
| Defocus Range (μm) | -0.5/-1.5 |
| Pixel Size (Å) | 0.8248 |
| Symmetry Imposed | C1 |
| Initial Particle Images | 2,475,999 |
| Final Particle Images | 601,617 |
| Map Resolution (Å) | 1.84 |
| FSC Threshold | 0.143 |

**Table S3. Model Refinement Statistics**

| Model component | |
|---|---|
| Model resolution (Å) | 1.95 |
| FSC threshold | 0.5 |
| Map sharpening *B* factor (Å$^2$) | -38.2 |
| Model composition | |
| Non-hydrogen atoms | 146561 |
| $Mg^{2+}$ ions | 284 |
| Waters | 6114 |
| Mean *B* Factors (Å$^2$) | |
| RNA | 24.25 |
| Protein | 24.91 |
| Waters | 16.95 |
| Other | 21.21 |
| R.m.s. deviations from ideal values | |
| Bond (Å) | 0.007 |
| Angle (°) | 0.968 |
| Molprobity score | 2.11 |
| Clash Score | 6.94 |
| Rotamer outliers (%) | 3.59 |
| Ramachandran plot | |
| Favored (%) | 95.61 |
| Allowed (%) | 3.73 |
| Outliers (%) | 0.66 |
| RNA validation | |
| Angles outliers (%) | 0.001 |
| Sugar pucker outliers (%) | 0.009 |
| Average suiteness | 0.589 |
